# FOLR2 Identifies Functionally Distinct Decidual Macrophage Populations that Coordinate Immune and Tissue Remodelling During Human Labour

**DOI:** 10.64898/2026.11.16.767055

**Authors:** Emilie O. Paterson, Katerina Rekopoulou, Annabel T.S. Chi, Shreya Sheth, Alice M. Buchan, Brendan Browne, Daniel J. Short, Lynne Sykes, Victoria Male

**Affiliations:** Department of Metabolism, Digestion and Reproduction, Imperial College London, UK; BRC Genomics Facility, Imperial College London, UK; Department of Obstetrics and Gynaecology, Chelsea and Westminster Hospital, London, UK; March of Dimes Prematurity Research Centre, Imperial College London, UK; The Parasol Foundation Centre for Women’s Health and Cancer Research, London, UK

**Keywords:** Decidua, pregnancy, labour, monocytes, macrophages, FOLR2

## Abstract

The initiation of human labour remains incompletely understood, although mounting evidence implicates decidual macrophages in this process. Despite this, macrophage heterogeneity remains poorly defined, with current macrophage classification systems, including M1/M2, failing to capture their biological diversity. Here, we investigate whether macrophage differentiation state, defined by expression of the tissue residency marker FOLR2, provides a more informative framework for characterising human decidual macrophages at term. RNA sequencing and flow cytometry identified FOLR2- and FOLR2+ macrophages as transcriptionally and phenotypically distinct populations. FOLR2⁻ macrophages displayed features consistent with recently recruited monocyte-derived cells and were characterised by enrichment of inflammatory and leukocyte recruitment pathways, with greater expression of IL-8 and CCL5 than FOLR2⁺ macrophages. In contrast, FOLR2⁺ macrophages expressed higher levels of MHC class II molecules, CCL2, IL-6, MMP9 and MMP19, and demonstrated the capacity to degrade collagen in fetal membrane explants. Monocyte and macrophage numbers peaked during early labour in the decidua parietalis, accompanied by increased CCL5 expression, whereas FOLR2+ macrophage expression of MMP9 was highest prior to labour. These findings suggest that decidual macrophage heterogeneity is best understood through differentiation state rather than M1/M2 polarisation and point to complementary roles for FOLR2⁻ and FOLR2⁺ macrophages in leukocyte recruitment, tissue remodelling and the local adaptations required for labour initiation and progression.

## Introduction

Labour is a highly coordinated process involving endocrine, inflammatory, and mechanical changes. While the feedback mechanisms that sustain labour after its initiation are well characterised, the precise molecular and cellular events that trigger its onset remain unknown. One longstanding hypothesis is that labour may be initiated by an immune signal originating in the uterine mucosa, called the decidua. Macrophages are a major leukocyte population in the decidua, comprising approximately 10–20% of decidual immune cells (1–4). Decidual macrophages have been reported to increase in frequency during human labour and are therefore attractive candidates as initiators of the putative immune signal that triggers labour (5–7). Although the logistics of sample collection make it difficult to distinguish cause from effect in humans, studies in animal models show that macrophage frequency increases before the onset of labour, supporting a potential role for these cells in labour initiation (7,8).

The decidua provides a unique physiological setting for investigating macrophage plasticity because the tissue environment shifts throughout pregnancy with a transient state that shares features with inflammation at implantation, a relatively quiescent phase in the second trimester, and a return to heightened inflammation as parturition approaches (9). Labour offers a rare opportunity to investigate macrophage functions in a rapidly changing tissue that may include both inflammatory, tissue remodelling and homeostatic aspects simultaneously. Notably, the decidua comprises sites with differing functions, creating distinct microenvironments that may shape local macrophage populations in different ways. The decidua basalis lies beneath the placenta and directly interacts with invading placental trophoblast cells while the decidua parietalis lines the uterine cavity and interacts with the fetal membranes (10,11).

Historically, macrophages have been categorised into two polarised states: the pro inflammatory M1 subtype and the anti-inflammatory M2 subtype, defined based on surface marker expression and cytokine production (12). However, these observations have not been validated *in vivo*, and several authors have highlighted the lack of evidence supporting discrete M1 and M2 identities in tissues (13–15). Rather, there is a growing consensus that macrophages exist along a spectrum of activation states adapted to the local tissue environment (16,17).

In first trimester decidua, where M1 and M2 phenotypes have been defined using CD80, CD163 and CD209, the majority of macrophages are phenotypically M2-like but produce both M1 and M2 cytokines (18). In third trimester decidua parietalis, the majority of macrophages have neither an M1 nor an M2 phenotype (19). CD11c has been used as a marker associated with M1 macrophages, based on studies of adipose tissue macrophages in mice and humans (20,21): this marker has subsequently been used to define macrophage subsets in first trimester human decidua, although neither subset conforms to classical M1 or M2 polarisation (4,22–25). Subsets distinguished by CD11c expression have also been identified in term decidua basalis but, in contrast to adipose tissue and first trimester decidua, in third trimester decidua it is the CD11c^hi^ population that more closely resembles an M2 phenotype (26). However, term CD11c^hi^ decidual macrophages also display surface markers typically linked to M1-like profiles, highlighting that neither of these subsets can be unambiguously classified as M1 or M2.

These findings suggest that the M1/M2 paradigm does not adequately capture the diversity of macrophage states in the decidua, highlighting the need for a more robust and biologically meaningful approach to decidual macrophage phenotyping. One possibility is to consider the spectrum of differentiation states of myeloid cells in the decidua. In this framework, incoming monocytes progressively acquire the macrophage marker CD68, followed by FOLR2, reflecting the establishment of tissue residency (27,28). This framework therefore allows distinction between recently recruited monocyte-derived cells and macrophages that have spent longer in the decidual microenvironment, with the recent development of drugs targeting myeloid cell recruitment (29,30) and FOLR2 (31,32) enhancing its translational relevance.

In this paper we characterise two phenotypically and functionally distinct populations of macrophages, distinguished by the tissue residency marker FOLR2, in the decidua at term. We go on to describe population frequencies and phenotype in the decidua basalis and parietalis before labour, and in early or established labour. Our findings lead us to propose that that differentiation state provides the most informative framework for characterising decidual macrophage heterogeneity and support a role for decidual macrophages in coordinating the local tissue adaptations required for labour initiation and progression.

## Materials and Methods

### Human tissue

Human samples used in this research project were obtained from the Imperial College Healthcare Tissue and Biobank (ICHTB). ICHTB is approved by Wales REC3 to release human material for research (22/WA/0214), and the samples for this project (R22010) were issued from sub-collections MET_VM_22_002, MET_VM_22_003 and MET_VM_23_001.

Placental tissues were collected from women delivering at term (>37 weeks’ gestation) and classified according to labour status at the time of delivery. Term not in labour (TNL) samples were obtained from women undergoing elective caesarean birth prior to the onset of labour and in the absence of regular uterine contractions. Term early labour (TEaL) samples were obtained from women scheduled for elective caesarean delivery who entered spontaneous labour prior to their planned procedure. These women presented with regular painful uterine contractions and cervical dilatation of less than 4 cm, consistent with the latent phase of labour. As tissue collection was only possible at delivery, all TEaL samples were obtained by caesarean delivery before progression to established labour.

Term labour (TL) samples were obtained following spontaneous vaginal delivery in established labour. All participants were considered to have low-risk pregnancies with no diagnosed obstetric complications or systemic disease. Donor characteristics are summarised in Supplementary Table 1.

### Dissection of decidua

Placentae were processed within 45 minutes of delivery. Placentae were dissected and immune cells isolated as described previously (33), with minor modifications, as described briefly. The maternal surface of the placenta was rinsed with PBS and blotted to remove excess blood. Using blunt forceps and disposable scalpels, thin sheets of decidua basalis were dissected and transferred into PBS-filled beakers. Tissues were stirred gently using a magnetic stirrer for 20 minutes to remove excess blood. Residual villous tissue was mechanically separated from the decidua using dissection scissors and a cell scraper. Cleaned decidua basalis tissue was transferred to 50 mL falcon tubes, suspended in PBS, and weighed.

Decidual membranes (amnion, chorion, and parietalis) were separated from the placenta using dissection scissors. The amnion was removed by peeling the membranes apart to improve visibility. Approximately 10 cm² sections of membrane were placed with the maternal side facing upwards on petri dishes, and the decidua parietalis was mechanically scraped using a sterile cell scraper. Scraped tissue was transferred to PBS and weighed.

The proximity between decidual tissue and placental villi makes complete exclusion of Hofbauer cells from dissected decidua basalis technically challenging and introduces the possibility that fetal-derived macrophages are present within populations isolated from the decidua. To determine the frequency of contaminating Hofbauer cells in decidua basalis macrophage preparations, we additionally stained for HLA-A2, HLA-A3 and HLA-B7, allowing the distinction between fetal-and maternal-derived cells where there is a mismatch in the expression of one of these HLA molecules. Among 13 decidua basalis preparations, Hofbauer cells were identified in two, with frequencies of 4.3% and 31.2% (Supplementary Figure 1).

### Immune cell isolation

Decidua was homogenised in GentleMACS C-tubes (Miltenyi Biotech) with Accutase (Invitrogen) at a 2:1 volume-to-tissue weight ratio. Samples were incubated (45 minutes, 37°C, 100 RPM). Digested tissue was filtered through 70 μm cell strainers and resuspended in RPMI medium containing penicillin/streptomycin. Cell suspensions were layered over Ficoll-Paque Plus (Cytiva) at a 2:1 ratio and centrifuged (400 x*g*, 20 minutes, RT, gentle acceleration and braking). The leukocyte-rich interphase was collected, washed in PBS, and resuspended for cell counting.

### Flow cytometry staining

Cells were stained immediately following isolation. All wash steps were carried out in 1 mL FACS buffer (apart from pre-viability stain where cells were washed in PBS), centrifuging at 500 x*g*, 5 mins, 4°C. Details of antibodies used are shown in Table 1.

**Table 1.** Antibodies used in ffow cytometry staining Flow sorting.

| Target | Fluorophore | Clone | Concentration | Supplier | RRID |
|---|---|---|---|---|---|
| **Surface – primary** |  |  |  |  |  |
| CCR2 | BV711 | K036C2 | 1/50 | Biolegend | AB_2800970 |
| CD11c | PE-Dazzle 594 | Bu15 | 1/400 | Biolegend | AB_2564548 |
| CD14 | BV510 | 63D3 | 1/25 | Biolegend | AB_2716229 |
| CD16 | BV605 | 3G8 | 1/200 | Biolegend | AB_2561354 |
| CD45 | APC-Cy7 | 2D1 | 1/50 | Biolegend | AB_2566376 |
| CD71 | FITC | CY1G4 | 1/25 | Biolegend | AB_2201482 |
| CD80 | BV785 | 2D10 | 1/100 | Biolegend | AB_2734271 |
| FOLR2 | APC | 94b/FOLR2 | 1/400 | Biolegend | AB_2721302 |
| HLA-A2 | BV785 | BB7.2 | 1/100 | Biolegend | AB_2721563 |
| HLA-A3 | FITC | GAP.A3 | 1/50 | Thermo Fisher | AB_2572503 |
| HLA-B7 | PE-Cy7 | BB7.1 | 1/100 | Thermo Fisher | AB_2802259 |
| HLA-DR | BV650 | L243 | 1/100 | Biolegend | AB_2562544 |
| HLA-DQ | PE | HLADQ1 | 1/100 | Biolegend | AB_604127 |
| MMP19 | Unconjugated | Polyclonal | 1/100 | Thermo Fisher | AB_2145660 |
| **Surface – secondary** |  |  |  |  |  |
| Rabbit IgG | AF680 | Polyclonal | 1/200 | Invitrogen | AB_2535758 |
| **Intracellular** |  |  |  |  |  |
| CCL2 | PE-Cy7 | 5D3-F7 | 1/200 | Biolegend | AB_2734490 |
| CCL5 | PerCP-Cy5.5 | VL1 | 1/25 | Biolegend | AB_2562665 |
| CCL18 | PE | REA487 | 1/200 | Miltenyi | AB_2904651 |
| CD68 | PerCP-Cy5.5 | Y1/82A | 1/25 | Biolegend | AB_10683171 |
| IL-6 | PE-Cy7 | MQ2-13A5 | 1/25 | Biolegend | AB_2572041 |
| IL-8 | AF700 | E8N1 | 1/25 | Biolegend | AB_2832819 |
| MMP9 | PE | EP1254 | 1/25 | abcam | AB_3754784 |

Following viability staining with fixable viability dye (eBioscience; 1/1000; 4°C, 15 minutes) cells were washed, surface staining was performed in FACS buffer (15 minutes, 4°C, in the dark) and cells were washed again. Where a secondary antibody was required to detect surface staining with unconjugated antibody, the secondary stain (15 minutes, 4°C, in the dark) was added following this wash. Cells were fixed (30 minutes, RT, in the dark) and finally washed and permeabilized using the Intracellular fixation and permeabilization buffer set (eBioscience) and stained intracellularly (30 minutes, RT, in the dark). The cells were then washed and resuspended in 400µL of FACS buffer for analysis. Samples were acquired on a Fortessa X20 cytometer using FACSDiva software (BD). Acquired results were analysed with FlowJo (Treestar).

### Flow sorting

Decidua parietalis immune cells were stained with fixable viability dye, CD45, CD14, CD71 and FOLR2. For sorting, macrophages were identified as live CD45+ CD14+ CD71+ cells. Within the macrophage gate, FOLR2+ and FOLR2-macrophage subsets were identified and sorted using the Sony SH800S directly into 15 mL falcon tubes containing 200 µL of culture medium.

### RNA sequencing

Cells were pelleted and lysed in RLT buffer (Qiagen) supplemented with 10% DMSO. The lysates were stored at minus 80 for later extraction of total RNA using the RNeasy Micro kit (Qiagen), according to the manufacturer’s instructions. RNA concentration and quality were assessed using an Agilent Tapestation: cDNA libraries were then prepared from 200 ng starting material using a NEBNext Single Cell/Low Input RNA Library Prep Kit (Illumina). Libraries were sequenced on a NextSeq 2000.

Alignments were generated using the nf-core/rnaseq 69 pipeline (34). Transcripts generated were adapter trimmed with Trim Galore! (v0.6.5) and mapped against the human reference genome (GRCh38.p14) using STAR (v2.7.11b) and Salmon (v.1.11.4). Length scaled gene counts generated by Salmon were used for downstream analysis using the iDEP platform (35). Length scaled gene counts imported and filtered to remove low-abundance transcripts. Principal component analysis (PCA) was performed as a quality control check, and to assess transcriptional separation between macrophage subsets. Differential gene expression analysis was conducted using DESeq2 with donor included as a blocking factor to account for matched sample structure. Differentially expressed genes were identified using a false discovery rate (FDR)-adjusted p-value threshold of <0.05 and a minimum fold-change threshold of 1.5.

To identify biological pathways associated with each macrophage subset, Gene Set Enrichment Analysis (GSEA) was performed using Gene Ontology (GO) Biological Process pathways. Pathways with FDR-adjusted p-values < 0.05 were considered significantly enriched.

### Term membrane incubation with decidual macrophage subsets

Placentae were collected from not-in-labour low-risk pregnancies. The chorion, amnion and decidua parietalis were separated from each other and immune cells isolated from the decidua parietalis as described above. After the removal of the decidua parietalis, from the remaining chorion, four uniform circular membrane pieces approximately 12mm across were excised using a scalpel.

Sorted macrophages were resuspended at 1×10^6^ cells/mL in RPMI plus 2% penicillin/streptomycin, non-essential amino acids, sodium pyruvate, 2-mercapto-ethanol and 1% FCS (all Gibco) and added to the membrane pieces. The negative control for membrane degradation was medium only and the positive control was media with the addition of 0.01% w/v collagenase IV (Life Technologies). Cells and membrane pieces were subsequently cultured for 24 hours (37°C, 5% CO2).

Following incubation, membrane pieces were removed and cut in half using a scalpel. Half of the cut membrane pieces were fixed in 10% neutral buffered formalin for 24 hours, then embedded in paraffin, sectioned and stained with Picrosirius red. Slides were imaged on an Olympus BX60F light microscope and analysed using the QuantSeg macro in FIGI (36). The other half of the membrane piece was snap frozen for later assessment of collagen content by hydroxyproline assay (Merck).

## Statistical analysis

Statistical testing and graph generation were performed using GraphPad Prism version 10.1.1. Data were assessed for normality using the Shapiro-Wilk and Kolmogorov-Smirnov tests. Statistical analyses were selected according to the normality of the data and experimental design (number of groups, repeated measures). When sphericity assumptions were not met, the Geisser Greenhouse correction was applied. Specific statistical tests used are described in the relevant figure legends.

## Results

### FOLR2 identifies two populations of macrophages cells in term decidua

We first sought to determine whether decidual myeloid cells could be resolved into distinct populations that phenotypically aligned with monocytes, recently recruited macrophages, and tissue-resident macrophages.

Placentae with attached fetal membranes and decidua were collected from women delivering by elective caesarean birth prior to the onset of labour at term. Immune cells were isolated from the decidua parietalis, adjoining the fetal membranes, and from the decidua basalis, attached to the placenta, and examined by flow cytometry. Myeloid lineage cells were identified as CD14+ and within this population, macrophages were distinguished from monocytes by their expression of CD68. FOLR2 staining of CD68+ macrophages consistently produced distinct FOLR2-and FOLR2+ populations (Figure 1A; Supplementary Figure 2A).

**Figure 1.**
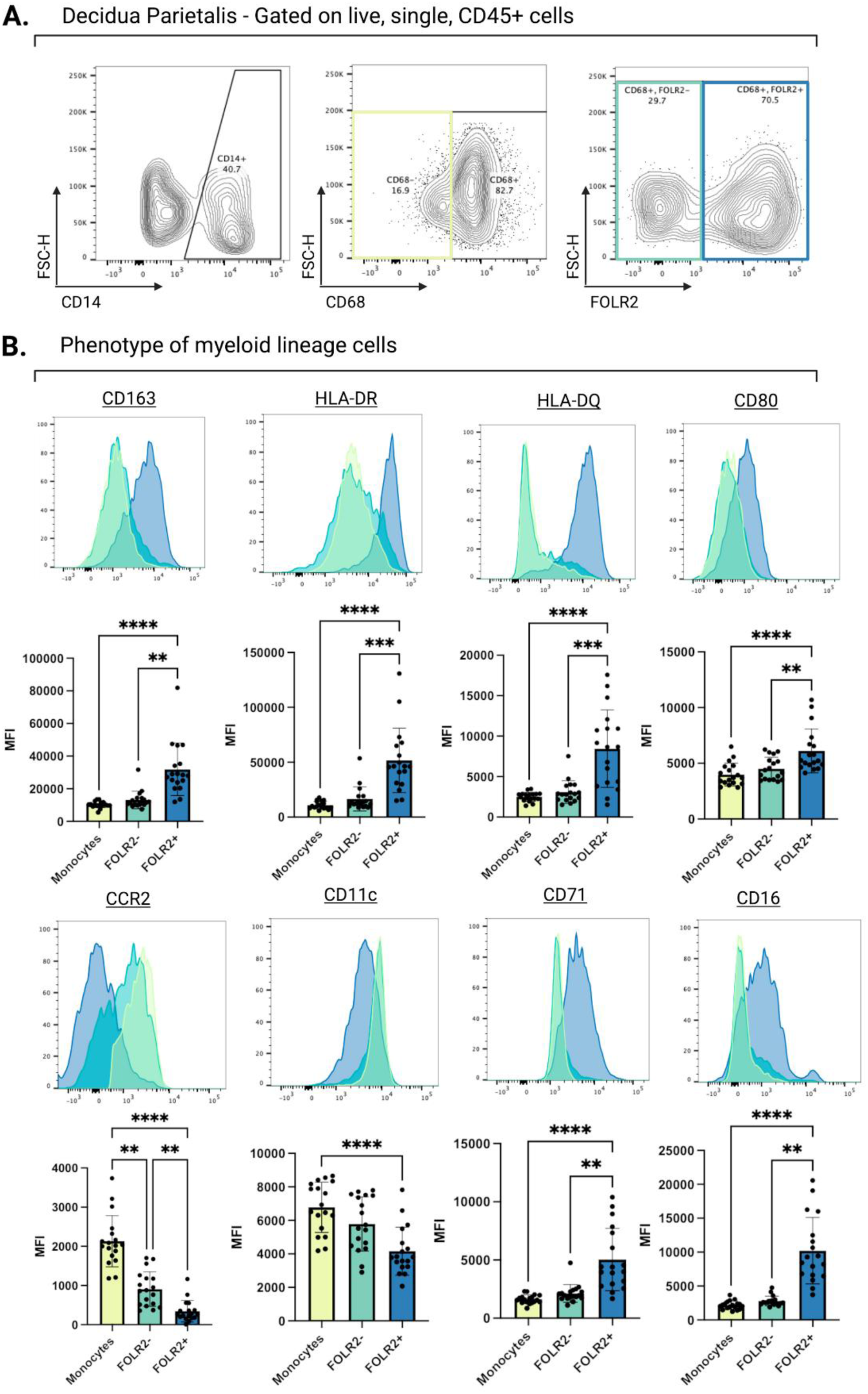
FOLR2 identifies two populations of macrophages cells in term decidua parietalis Placentae with attached fetal membranes and decidua were collected from women delivering by elective caesarean birth prior to the onset of labour at term. Immune cells were isolated from the decidua parietalis and examined by flow cytometry. Immune cells were identified as live, single CD45+ cells. **A.** Myeloid lineage cells were identified as CD14+. Monocytes (yellow gate) were defined as CD68-and macrophages as CD68+. Within the macrophage population, FOLR2-(green gate) and FOLR2+ (blue gate) subsets were defined. **B.** Expression of myeloid lineage markers by monocytes (yellow), FOLR2-(green) and FOLR2+ (blue) macrophages. Representative histograms and complete summary data for each marker examined (n = 18) are shown. Statistical comparisons were performed with Friedman’s test and Dunn’s post-hoc test. * p < 0.05; ** p < 0.01; *** p < 0.001; **** p < 0.0001.

Monocytes and FOLR2-macrophages were phenotypically similar, with no statistically significant difference in their expression of eight macrophage markers examined (CD163, HLA-DR, HLA-DQ, CD80, CCR2, CD11c, CD71, CD16; Figure 1B; Supplementary Figure 2B). Except for CD11c and CCR2, all these markers were more highly expressed in FOLR2+ macrophages than both monocytes and FOLR2-macrophages. CCR2 and CD11c were most highly expressed in monocytes, with FOLR2-macrophages expressing intermediate levels of these markers.

### FOLR2-negative and-positive macrophages are transcriptionally distinct

To confirm that FOLR2 expression identifies distinct subsets of decidual macrophages, RNA sequencing was performed on FOLR2+ and FOLR2-macrophages flow sorted from decidua parietalis collected from women delivering by elective caesarean birth prior to the onset of labour at term.

Principal component analysis demonstrated the separation of FOLR2+ and FOLR2-macrophages along PC1, which accounted for 28.9% of variation, indicating that these populations represent distinct macrophage subsets (Figure 2A). Differential gene expression analysis further identified marked differences in transcriptional profiles between the two populations (Figure 2B). As expected, FOLR2+ macrophages expressed higher levels of FOLR2, while FOLR2-macrophages expressed higher levels of genes associated with monocytes (S100A12, FCN1).

**Figure 2.**
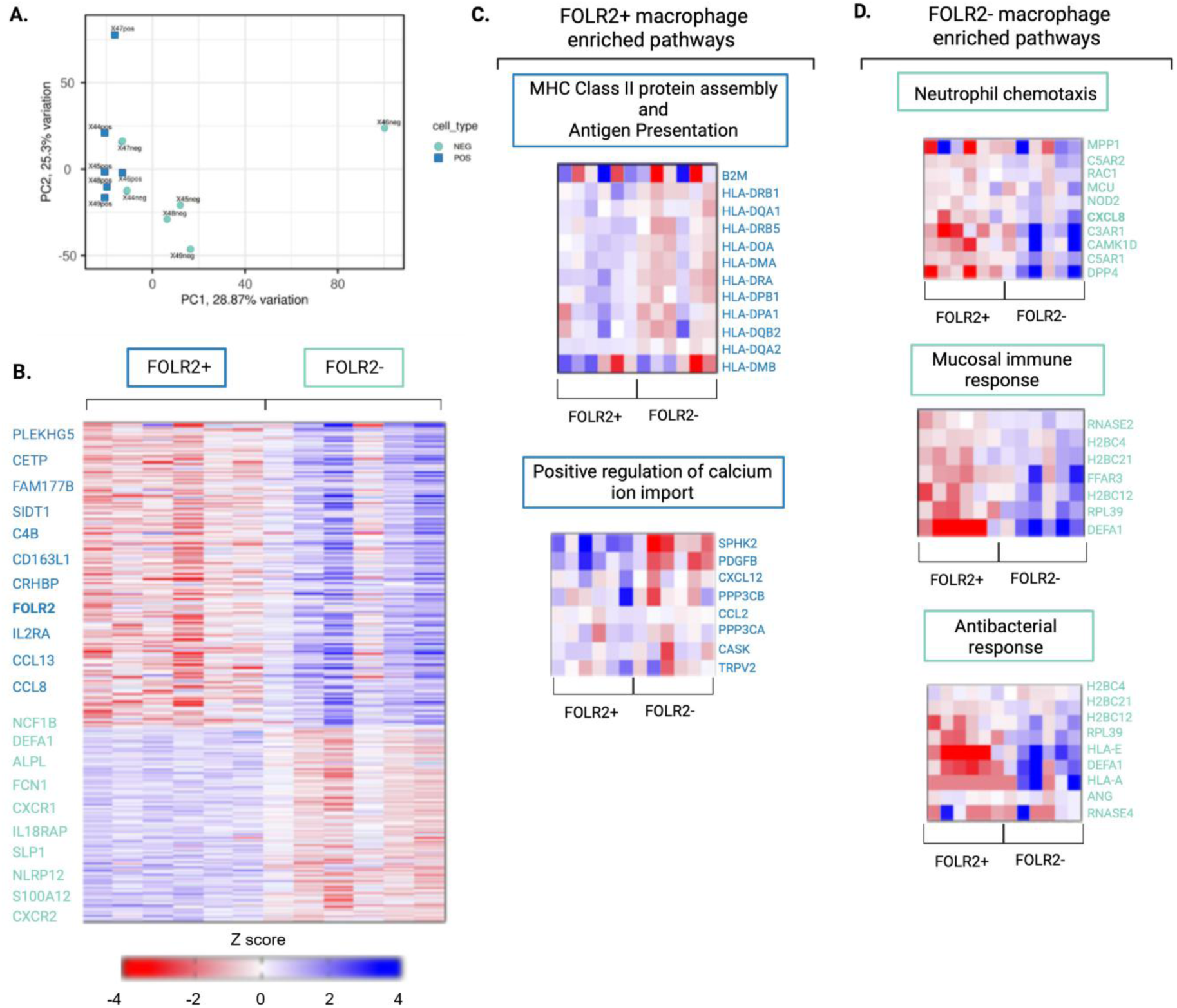
FOLR2-negative and-positive macrophages are transcriptionally distinct RNA sequencing was performed on FOLR2+ and FOLR2-macrophages flow sorted from term decidua parietalis (n = 6 of each). **A.** Principal component analysis. **B.** Differential gene expression analysis. Genes that are differentially expressed at FC > 1.5 and FDR-adjusted p < 0.05 are shown. **C,D.** Gene Ontology (GO) biological process gene sets enriched in FOLR2+ (**C**) and FOLR2-(**D**) decidual macrophages. Pathways with FDR-adjusted p < 0.05 are shown.

Gene set enrichment analysis identified enrichment of pathways associated with antigen processing and presentation and regulation of calcium ion import in FOLR2+ macrophages (Figure 2C), with the overexpression of genes encoding MHC class II molecules consistent with the increased expression of these molecules we observed at the protein level (Figure 1B). Meanwhile, FOLR2-macrophages demonstrated enrichment of pathways associated with neutrophil chemotaxis, mucosal immune responses and antibacterial responses (Figure 2D).

### FOLR2-negative and-positive macrophages have distinct cytokine production profiles

To investigate cellular function, we assessed cytokine and chemokine expression by decidual monocyte and macrophage subsets directly *ex vivo*. The selection of cytokines and chemokines to be examined was informed by their established roles in immune cell recruitment and inflammatory signalling. CCL2 is a well-established monocyte chemoattractant, while IL-8 (CXCL8), although classically associated with neutrophil recruitment, has also been shown to contribute to monocyte chemotaxis in human decidual cell-conditioned medium (37). CCL5 acts more broadly as a leukocyte chemoattractant, with implications of recruiting monocytes and macrophages to the decidua (37,38) and CCL18 has been associated with lymphocyte recruitment and macrophage differentiation (39,40). IL-6 was additionally included as previous work showed decidual macrophages producing it during late gestation and labour (41).

Overall, clear differences in cytokine expression were observed between these monocytes and the two macrophage populations, supporting their classification as distinct subsets. In line with findings at the transcriptional level (Figure 2C,D), a greater proportion of FOLR2-macrophages expressed IL-8 (Figure 3A) whereas FOLR2+ macrophages expressed high levels of CCL2 (Figure 3D). Both macrophage subsets expressed significantly more CCL5 than monocytes (Figure 3C). IL-6 was expressed by the majority of FOLR2+ macrophages, but not by monocytes, with only a minority of FOLR2-macrophages expressing IL-6, intermediate between the two other populations (Figure 3B). In contrast, CCL18 was most highly expressed by monocytes and FOLR2-macrophages, with FOLR2+ macrophages expressing lower levels (Figure 3E).

**Figure 3.**
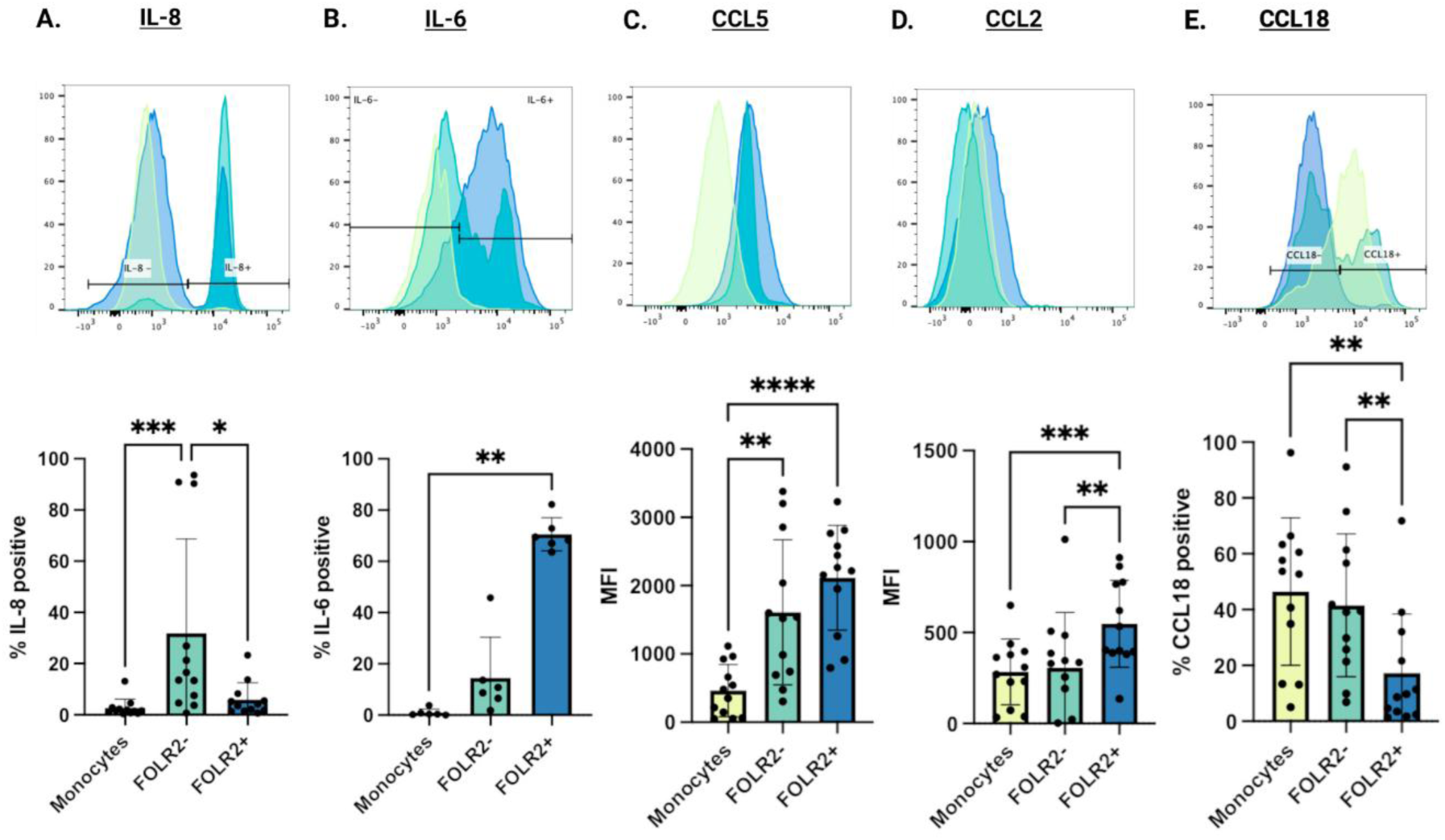
FOLR2-negative and-positive macrophages have distinct cytokine production profiles Decidual monocyte and macrophage populations freshly isolated from term decidua parietalis as shown in Figure 1 were stained intracellularly to examine their constitutive production of IL-8 (**A**, n = 12), IL-6 (**B**, n = 6), CCL5 (**C**, n = 12), CCL2 (**D**, n = 12) and CCL18 (**E**, n = 12). Representative histograms and complete summary data for cytokine production by monocytes (yellow), FOLR2-(green) and FOLR2+ (blue) macrophages are shown. Statistical comparisons were performed with Friedman’s test and Dunn’s post-hoc test (non-normally distributed data: IL-8, IL-6, CCL18) and repeated measures one-way ANOVA with Tukey’s multiple comparisons test (normally distributed data: CCL5, CCL2). * p < 0.05; ** p < 0.01; *** p < 0.001; **** p < 0.0001.

### FOLR2-positive macrophages express MMPs and can degrade fetal membrane collagen

Decidual macrophages have been reported to express matrix metalloproteinases (MMPs) which degrade structural matrix proteins including collagen, elastin, fibronectin, and microfibrils and could therefore contribute to fetal membrane weakening and rupture (42,43). Therefore, we characterised MMP expression by decidual macrophage subsets, finding that monocytes, FOLR2-and FOLR2+ decidual macrophages all express some MMP9 and MMP19, but that FOLR2+ decidual macrophages do so at a significantly higher level than the other subsets (Figure 4A,B).

**Figure 4.**
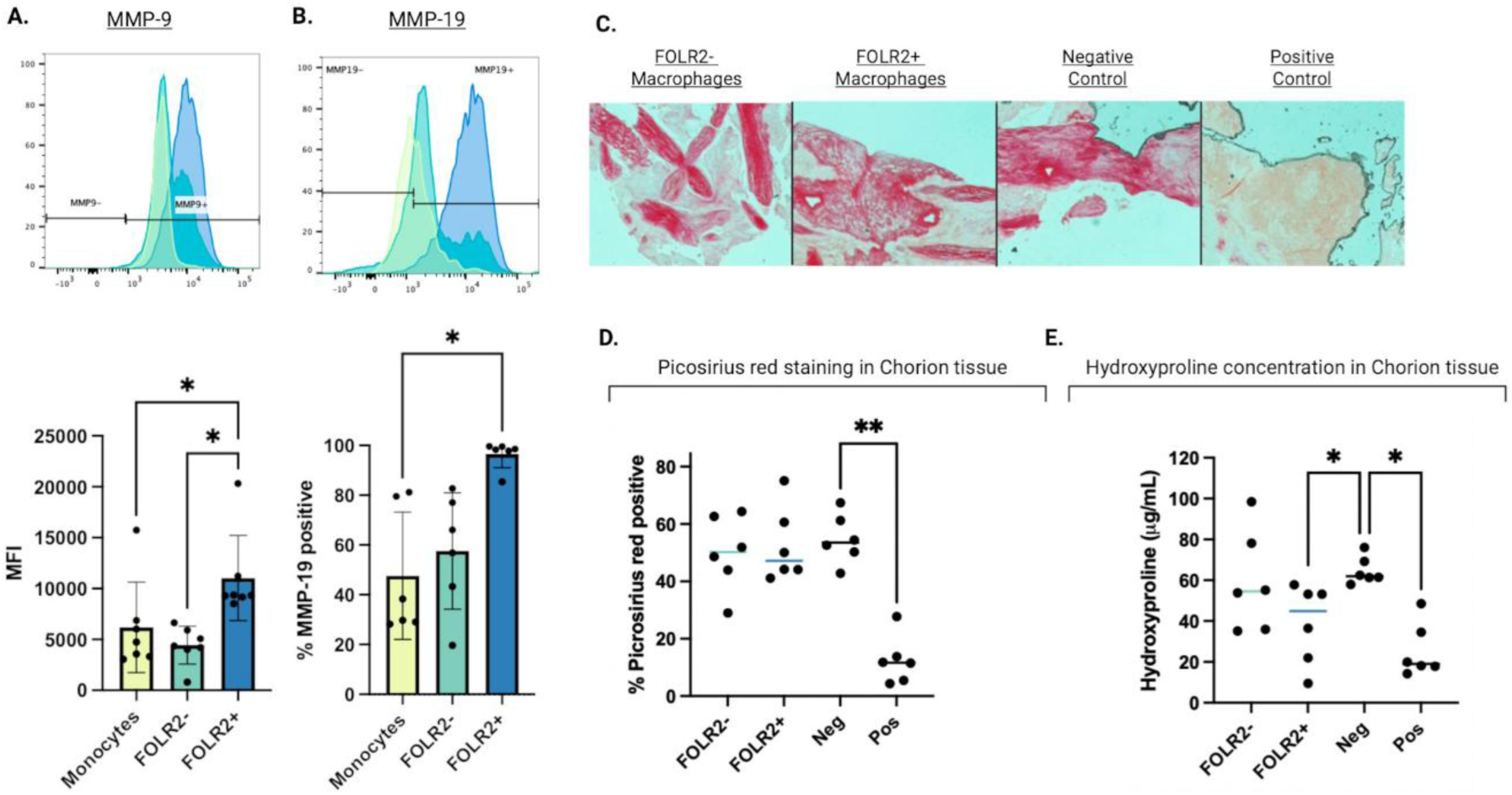
FOLR2-positive macrophages express MMPs and can degrade fetal membrane collagen **A,B.** Decidual monocyte and macrophage populations freshly isolated from term decidua parietalis as shown in Figure 1 were stained intracellularly for MMP9 (**A**, n = 7) and surfaced stained for MMP19 (**B**, n = 6). Representative histograms and complete summary data for monocytes (yellow), FOLR2-(green) and FOLR2+ (blue) macrophages are shown. Statistical comparisons were performed with Friedman’s test and Dunn’s post-hoc test, * p < 0.05. **C-E.** FOLR2- and FOLR2+ macrophages were flow sorted from term decidua parietalis and cultured with isolated chorion membrane tissue for 24 h. The positive control was collagenase IV treatment and the negative control was culture in medium alone. Remaining collagen content was assessed by Picrosirus red staining and hydroxyproline quantification. Representative images (**C**) and summary data (**D, E**, n = 6 per group) are shown. Statistical comparisons were performed with repeated measures one-way ANOVA with Tukey’s multiple comparisons test (**D**, data normally distributed) or Friedman’s test and Dunn’s post-hoc test (**E**, data non-normally distributed). * p < 0.05; ** p < 0.01.

To assess the functional significance of these differences, we next investigated the ability of decidual macrophage subsets to contribute to extracellular matrix degradation of fetal membranes. FOLR2- and FOLR2+ macrophages were flow sorted from decidua parietalis collected from women delivering by elective caesarean birth prior to the onset of labour at term and cultured with isolated chorion membrane tissue for 24 h. Remaining collagen content was assessed using Picrosirus red staining (Figure 4C, D) and hydroxyproline quantification (Figure 4E). Collagenase IV treatment was used as the positive control and culture in medium only was used as the negative control.

As expected, collagenase treatment led to significant degradation of fetal membrane collagen (Figure 4D,E). In line with our finding that FOLR2+ macrophages express more MMP9 and-19 than FOLR2-macrophages, fetal membrane co-culture with FOLR2+ macrophages was associated with a reduction in collagen content, although this was only significant when measured by hydroxyproline assay, which is more sensitive and also more robust to by tissue sampling, collagen organisation and image-analysis parameters.

### Monocytes and macrophages change in number and activity over the course of labour

Given the proposed role for macrophages in the initiation of labour, we next sought to define how the number and activity of monocytes and the two macrophage subsets change in early and established labour. Term not in labour (TNL) samples were collected from women delivering by elective caesarean birth prior to the onset of labour; term early labour (TEaL) samples were collected from women scheduled for caesarean delivery who entered spontaneous labour prior to their scheduled procedure but were delivered by caesarean before reaching 4cm of cervical dilatation; term labour (TL) samples were collected following spontaneous vaginal delivery, having been in established labour.

In the decidua parietalis, the absolute number of monocytes, FOLR2-macrophages and FOLR2+ macrophages per gram of tissue was increased in early labour, compared to either not in labour or following established labour, although the difference was only statistically significant for monocytes (Figure 5A). Interestingly, this pattern was not present in the decidua basalis, in which the absolute number of all three cell types remained constant across labour states (Figure 5B).

**Figure 5.**
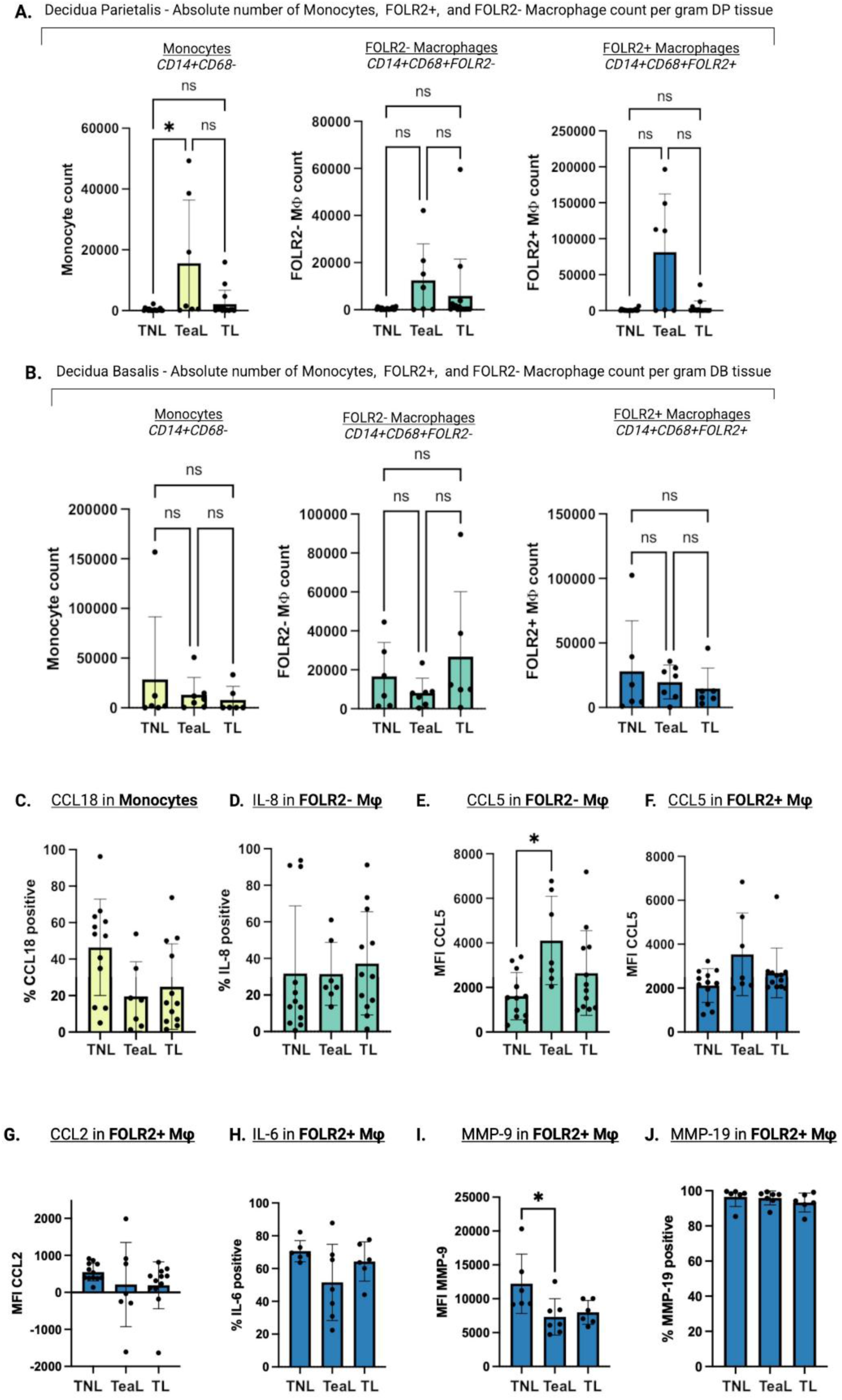
Monocytes and macrophages change in number and activity over the course of labour **A,B.** Immune cells were isolated from decidua parietalis (**A**) or basalis (**B**) and monocyte and macrophage populations identified by flow cytometry as shown in Figure 1. Absolute numbers of cells per gram of tissue in samples collected at different labour statuses are shown (decidua parietalis TNL, n = 18; TEaL, n = 7; TL, n = 12; decidua parietalis, n = 6 of each). **C**-**H.** Cells were stained intracellularly to examine their constitutive production of CCL18 (**C**), IL-8 (**D**), CCL5 (**E,F**), CCL2 (**G**) and IL-6 (**H**). Production across labour statuses in indicated cell subsets is shown. **I,J.** Cells were stained intracellularly for MMP9 (**I**) and surfaced stained for MMP19 (**J**). Statistical comparisons were performed with Kruskal Wallis with Dunn’s post-hoc test. * p < 0.05. TNL, term not in labour; TEaL, term early labour; TL term labour.

We next examined how the cytokines produced by each subset changed depending on labour status. The major cytokines produced by each subset (as identified in Figure 3) are shown in Figure 5C-H: complete data are shown in Supplementary Figure 3. The major chemokine produced by monocytes in the decidua parietalis, CCL18, tended to be produced by a higher proportion of cells in non-labouring samples, compared to early or established labour, although the difference was not statistically significant (Figure 5C). The proportion of FOLR2-macrophages expressing IL-8, a chemokine characteristic of this subset, did not change in these cells by labour status (Figure 5D). CCL5, which was made by both FOLR2- and FOLR2+ macrophages, was maximally produced by both subsets during early labour, although this was only significant for FOLR2-macrophages (Figure 5E,F). CCL2 and IL-6 were mostly produced by FOLR2+ macrophages, but their production did not differ significantly by labour status (Figure 5G,H).

Finally, we examined the production of MMPs by FOLR2+ macrophages, since this subset was the major source of MMPs, and was able to degrade fetal membrane collagen. Interestingly, we found significantly higher expression of MMP9 before the initiation of labour (Figure 5I). There was no significant difference in the proportion of FOLR2+ macrophages expressing MMP19 by labour status (Figure 5J). Expression of MMP9 and-19 did not differ by labour status in monocytes or FOLR2-macrophages (Supplementary Figure 4).

## Discussion

A major challenge in the study of decidual macrophages has been the lack of a phenotyping strategy that can be applied consistently across gestation, anatomical location and labour status. In particular, the M1/M2 paradigm has proved insufficient to capture the complexity of tissue macrophage populations *in vivo* (13–15). Here, we establish that the tissue residency marker FOLR2 distinguishes two populations of decidual macrophages in both decidua basalis and parietalis at term, across labour statuses. The two subsets are transcriptomically and phenotypically distinct, with FOLR2-macrophages expressing higher levels of transcripts and proteins, notably IL-8 (*CXCL8*) and CCL5, associated with leukocyte recruitment and mucosal immune responses. FOLR2+ macrophages also expressed high levels of CCL5 but were distinguished from FOLR2-macrophages by their increased expression of MHC class II, CCL2, IL-6 and MMPs. FOLR2+ macrophages were also able to degrade collagen in explanted fetal membranes *in vitro*.

The higher proportion of cells expressing the neutrophil chemotactic protein IL-8, as well as the higher expression of the general leukocyte chemoattractant CCL5, by FOLR2-macrophages is consistent with a broadly pro-inflammatory role. The FOLR2+ subset also expresses CCL5, which is widely recognised as a pro-inflammatory cytokine. CCL2 and IL-6 are also usually considered to be pro-inflammatory cytokines, but these also have roles in regulated tissue remodelling and homeostasis, depending on the other cytokines present in the tissue and, in the case of IL-6, the availability of soluble and membrane-bound receptors (44,45). These contextual aspects are still to be defined in the decidua. The expression of FOLR2 is most often considered to be associated with a tissue-resident phenotype (27,28) and that of MMP9 and MMP19 indicates a role in extracellular matrix and tissue remodelling. Our finding that FOLR2+ macrophages can degrade collagen in fetal membranes potentially suggests a role in the rupture of the membranes, a key event in the initiation of labour.

These findings are consistent with a recent study which identified two major macrophage subsets in term decidua basalis, distinguished by differential expression of HLA-DR (26). An HLA-DR^low^ subset expresses CCR2 and IL-8 and is therefore likely to align with FOLR2-decidual macrophages, while an HLA-DR^hi^ subset exhibits increased expression of FOLR2, CD163 and CD16 and is therefore likely to align with the FOLR2+ decidual macrophages. In line with our findings, the HLA-DR^low^ subset had a transcriptional profile associated with inflammation while the transcriptional profile of the HLA-DR^hi^ subset could not be unambiguously classified as either pro-inflammatory or tissue-homeostatic. Similarly, recent findings from a study in which FOLR2 was knocked out in the macrophage cell line THP-1 suggest that FOLR2 may have functions extending beyond its association with tissue homeostasis, including a potential role in inflammasome activation (46).

Further to work that has identified FOLR2 as a marker of tissue-resident macrophages, it has been proposed that recruited monocytes can contribute to the maintenance of tissue-resident macrophage populations (17,27,28). Our findings are consistent with monocytes recruited to the decidua differentiating into FOLR2-macrophages, and then into FOLR2+ macrophages, although importantly we have not formally demonstrated this occurring. FOLR2-decidual macrophages are phenotypically similar to monocytes and express some transcripts (S100A12, FCN1) that are characteristic of monocytes, consistent with FOLR2-macrophages having recently differentiated from recruited monocytes. Our identification of FOLR2-macrophages as expressing certain phenotypic markers (CD80, CCR2, CD11c) and cytokines (CCL2, CCL5, CCL18) at levels between monocytes and FOLR2+ macrophages supports the idea that this population represents an intermediate differentiation state. This suggestion is further supported by the recent report on macrophage subsets in the decidua basalis, defined by their expression of HLA-DR (26). This study used pseudotime inference of RNAseq data to identify a trajectory of differentiation starting with HLA-DR^low^ macrophages, which align with FOLR2-macrophages, and terminating in an HLA-DR^hi^ FOLR2+ population.

Our finding that the number of monocytes, FOLR2- and FOLR2+ macrophages in the decidua parietalis all peak during early labour is consistent with the differentiation of recently recruited monocytes to FOLR2- and FOLR2+ macrophages and also indicates that this process is involved in early labour, and potentially even the initiation of labour. That we did not observe similar labour-associated changes in the decidua basalis could suggest that the parietalis is the more immunologically active during labour, in line with the suggestion that the interface between the parietalis and chorion is the site of labour initiation (47,48). However, we did find that some decidua basalis samples included FOLR2+ macrophages of fetal origin, likely to represent Hofbauer cells from closely apposed placental villi. Since the frequency of Hofbauer cells is expected to remain relatively stable across labour, their presence could potentially mask labour-associated fluctuations within the FOLR2+ macrophage population in decidua basalis.

Expression of CCL5 by both macrophage populations also peaked during early labour, which could suggest feedback to recruit more monocytes, in line with their increased numbers at this time. MMP9 was most highly expressed by FOLR2+ macrophages in the decidua parietalis prior to labour, which potentially indicates that their function in tissue remodelling and weakening the fetal membranes primarily occurs before labour, again suggestive of a role in the initiation of labour. In support of this interpretation, FOLR2+ macrophages were able to degrade collagen in fetal membrane explants.

In addition to the limitations already discussed, several other important constraints of our approach should be considered in interpreting our findings. Comparisons between labour groups were necessarily cross-sectional, as longitudinal sampling of the decidua throughout labour is not possible, meaning subtle labour-associated changes may be obscured by biological variation between tissue donors. Where we identified trends that did not reach statistical significance, this may reflect limitations in statistical power associated with sample size rather than absence of biological relevance. However, the inclusion of clinically distinct labour groups, particularly the term early labour cohort, was a strength of the study, enabling assessment of transitional immune changes occurring during the progression from non-labouring to established labour states.

Our findings lead us to propose that macrophage heterogeneity in the decidua is better characterised by differentiation state and tissue residency than by the M1/M2 framework. We suggest that monocytes are recruited into the decidua during early labour, possibly under the influence of CCL2 produced by FOLR2+ macrophages and subsequently differentiate into FOLR2-macrophages. The majority of FOLR2-macrophages constitutively express IL-8, as well as increased expression of CCL5 during early labour, which may feed back to recruit further monocytes. FOLR2-macrophages give rise to FOLR2+ macrophages, the majority of which constitutively express IL-6 and MMPs, with MMP-9 expression highest prior to labour onset (Figure 6). Taken together, these findings suggest that FOLR2-macrophages may contribute primarily to leukocyte recruitment, whereas FOLR2+ macrophages may additionally contribute to tissue homeostasis and extracellular matrix remodelling, including weakening the fetal membranes. Through these complementary functions, decidual macrophage subsets may coordinate the immune and structural adaptations required for labour initiation and progression. Recently-developed drugs targeting myeloid cell recruitment (29,30) and FOLR2 (31,32) could modulate these functions, making decidual macrophages an attractive therapeutic target for the prevention or induction of labour.

**Figure 6.**
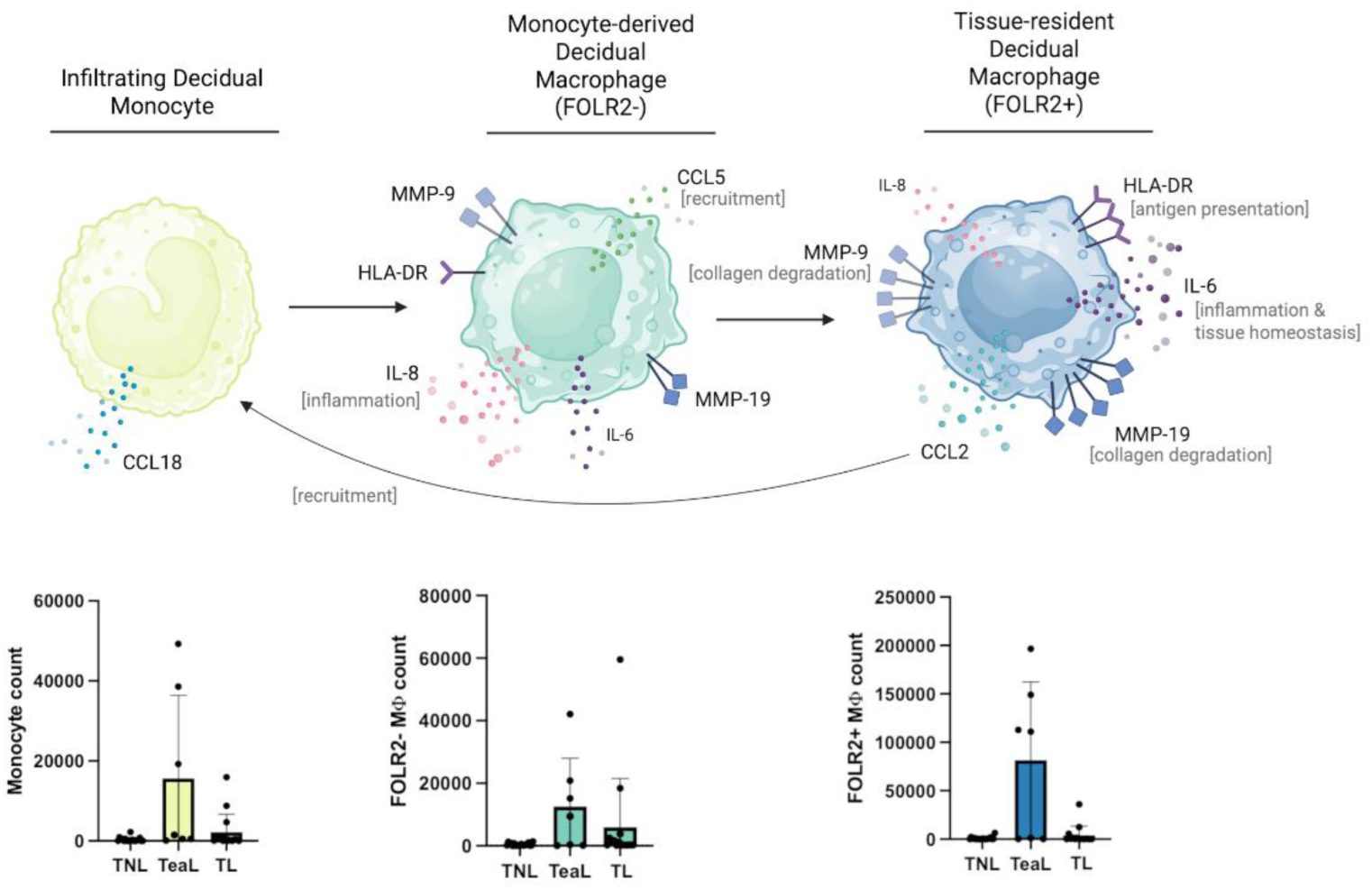
*Proposed model of decidual monocyte to macrophage differentiation and function in labour.* TNL, term not in labour; TEaL, term early labour; TL term labour. Created with BioRender.com.

## Data availability statement

The datasets presented in this study can be found in online repositories. The names of the repository/repositories and accession number(s) can be found below:

https://doi.org/10.17605/OSF.IO/JEK98

https://www.ncbi.nlm.nih.gov/geo/query/acc.cgi?acc=GSE337384

## Ethics statement

Human samples used in this research project were obtained from the Imperial College

Healthcare Tissue and Biobank (ICHTB). ICHTB is approved by Wales REC3 to release human material for research (22/WA/0214), and the samples for this project (R22010) were issued from sub-collections MET_VM_22_002, MET_VM_22_003 and MET_VM_23_001.

## Author contributions

Conceptualisation: EOP, VM. Investigation (performing experiments) and data curation: EOP, KR. Formal analysis: EOP, KR, AC, VM. Resources (identification and consent of tissue donors): EOP, SS, AB, BB, DS. Supervision: VM, LS. Funding acquisition: VM. Writing – original draft: EOP, VM. Writing – review and editing: all authors.

## Acknowledgements

We would like to thank patients at Chelsea and Westminster Hospital who contributed samples to this study. Samples collected with the support of the Imperial College Healthcare Tissue and Biobank (ICHTB) funded by the National Institute for Health Research (NIHR) Biomedical Research Centre based at Imperial College Healthcare NHS Trust and Imperial College London. The views expressed are those of the author(s) and not necessarily those of the NHS, the NIHR or the Department of Health.

## References

1. Bulmer JN, Johnson PM. Macrophage populations in the human placenta and amniochorion. Clin Exp Immunol. 1984 Aug;57(2):393–403. PubMed PMID: 6380834; PubMed Central PMCID: PMC1536121.

2. Rieger L, Honig A, Sütterlin M, Kapp M, Dietl J, Ruck P, et al. Antigen-presenting cells in human endometrium during the menstrual cycle compared to early pregnancy. J Soc Gynecol Investig. 2004 Oct;11(7):488–93. doi:10.1016/j.jsgi.2004.05.007 PubMed PMID: 15458747.

3. Bartmann C, Segerer SE, Rieger L, Kapp M, Sütterlin M, Kämmerer U. Quantification of the predominant immune cell populations in decidua throughout human pregnancy. Am J Reprod Immunol. 2014 Feb;71(2):109–19. doi:10.1111/aji.12185 PubMed PMID: 24330065.

4. Vento-Tormo R, Efremova M, Botting RA, Turco MY, Vento-Tormo M, Meyer KB, et al. Single-cell reconstruction of the early maternal–fetal interface in humans. Nature. 2018 Nov 15;563(7731):347–53. doi:10.1038/s41586-018-0698-6

5. Osman I. Leukocyte density and pro-inflammatory cytokine expression in human fetal membranes, decidua, cervix and myometrium before and during labour at term. Molecular Human Reproduction. 2003 Jan 1;9(1):41–5. doi:10.1093/molehr/gag001

6. Gomez-Lopez N, Vadillo-Perez L, Hernandez-Carbajal A, Godines-Enriquez M, Olson DM, Vadillo-Ortega F. Specific inflammatory microenvironments in the zones of the fetal membranes at term delivery. Am J Obstet Gynecol. 2011 Sep;205(3):235.e15-24. doi:10.1016/j.ajog.2011.04.019 PubMed PMID: 21763637.

7. Hamilton S, Oomomian Y, Stephen G, Shynlova O, Tower CL, Garrod A, et al. Macrophages Infiltrate the Human and Rat Decidua During Term and Preterm Labor: Evidence That Decidual Inflammation Precedes Labor1. Biology of Reproduction. 2012 Feb 1;86(2). doi:10.1095/biolreprod.111.095505

8. Shynlova O, Nedd-Roderique T, Li Y, Dorogin A, Nguyen T, Lye SJ. Infiltration of myeloid cells into decidua is a critical early event in the labour cascade and post-partum uterine remodelling. J Cellular Molecular Medi. 2013 Feb;17(2):311–24. doi:10.1111/jcmm.12012

9. Mor G, Cardenas I, Abrahams V, Guller S. Inflammation and pregnancy: the role of the immune system at the implantation site. Ann N Y Acad Sci. 2011 Mar;1221(1):80–7. doi:10.1111/j.1749-6632.2010.05938.x PubMed PMID: 21401634; PubMed Central PMCID: PMC3078586.

10. Yang F, Zheng Q, Jin L. Dynamic Function and Composition Changes of Immune Cells During Normal and Pathological Pregnancy at the Maternal-Fetal Interface. Front Immunol. 2019;10:2317. doi:10.3389/fimmu.2019.02317 PubMed PMID: 31681264; PubMed Central PMCID: PMC6813251.

11. Krop J, Tian X, van der Hoorn ML, Eikmans M. The Mac Is Back: The Role of Macrophages in Human Healthy and Complicated Pregnancies. Int J Mol Sci. 2023 Mar 10;24(6):5300. doi:10.3390/ijms24065300 PubMed PMID: 36982375; PubMed Central PMCID: PMC10049527.

12. Wynn TA, Chawla A, Pollard JW. Macrophage biology in development, homeostasis and disease. Nature. 2013 Apr 25;496(7446):445–55. doi:10.1038/nature12034 PubMed PMID: 23619691; PubMed Central PMCID: PMC3725458.

13. Italiani P, Boraschi D. From Monocytes to M1/M2 Macrophages: Phenotypical vs. Functional Differentiation. Front Immunol. 2014;5:514. doi:10.3389/fimmu.2014.00514 PubMed PMID: 25368618; PubMed Central PMCID: PMC4201108.

14. Mills CD, Ley K. M1 and M2 macrophages: the chicken and the egg of immunity. J Innate Immun. 2014;6(6):716–26. doi:10.1159/000364945 PubMed PMID: 25138714; PubMed Central PMCID: PMC4429858.

15. Strizova Z, Benesova I, Bartolini R, Novysedlak R, Cecrdlova E, Foley LK, et al. M1/M2 macrophages and their overlaps - myth or reality? Clin Sci (Lond). 2023 Aug 14;137(15):1067–93. doi:10.1042/CS20220531 PubMed PMID: 37530555; PubMed Central PMCID: PMC10407193.

16. Mosser DM, Edwards JP. Exploring the full spectrum of macrophage activation. Nat Rev Immunol. 2008 Dec;8(12):958–69. doi:10.1038/nri2448 PubMed PMID: 19029990; PubMed Central PMCID: PMC2724991.

17. Guilliams M, Thierry GR, Bonnardel J, Bajenoff M. Establishment and Maintenance of the Macrophage Niche. Immunity. 2020 Mar 17;52(3):434–51. doi:10.1016/j.immuni.2020.02.015 PubMed PMID: 32187515.

18. Svensson J, Jenmalm MC, Matussek A, Geffers R, Berg G, Ernerudh J. Macrophages at the Fetal–Maternal Interface Express Markers of Alternative Activation and Are Induced by M-CSF and IL-10. The Journal of Immunology. 2011 Oct 1;187(7):3671–82. doi:10.4049/jimmunol.1100130

19. Xu Y, Romero R, Miller D, Kadam L, Mial TN, Plazyo O, et al. An M1-like Macrophage Polarization in Decidual Tissue during Spontaneous Preterm Labor That Is Attenuated by Rosiglitazone Treatment. The Journal of Immunology. 2016 Mar 15;196(6):2476–91. doi:10.4049/jimmunol.1502055

20. Lumeng CN, Bodzin JL, Saltiel AR. Obesity induces a phenotypic switch in adipose tissue macrophage polarization. J Clin Invest. 2007 Jan;117(1):175–84. doi:10.1172/JCI29881 PubMed PMID: 17200717; PubMed Central PMCID: PMC1716210.

21. Hirata Y, Tabata M, Kurobe H, Motoki T, Akaike M, Nishio C, et al. Coronary atherosclerosis is associated with macrophage polarization in epicardial adipose tissue. J Am Coll Cardiol. 2011 Jul 12;58(3):248–55. doi:10.1016/j.jacc.2011.01.048 PubMed PMID: 21737014.

22. Houser BL, Tilburgs T, Hill J, Nicotra ML, Strominger JL. Two unique human decidual macrophage populations. J Immunol. 2011 Feb 15;186(4):2633–42. doi:10.4049/jimmunol.1003153 PubMed PMID: 21257965; PubMed Central PMCID: PMC3712354.

23. Jiang X, Du MR, Li M, Wang H. Three macrophage subsets are identified in the uterus during early human pregnancy. Cell Mol Immunol. 2018 Dec;15(12):1027–37. doi:10.1038/s41423-018-0008-0 PubMed PMID: 29618777; PubMed Central PMCID: PMC6269440.

24. Vondra S, Höbler AL, Lackner AI, Raffetseder J, Mihalic ZN, Vogel A, et al. The human placenta shapes the phenotype of decidual macrophages. Cell Rep. 2023 Jan 31;42(1):111977. doi:10.1016/j.celrep.2022.111977 PubMed PMID: 36640334.

25. Li Y, Sang Y, Chang Y, Xu C, Lin Y, Zhang Y, et al. A Galectin-9-Driven CD11chigh Decidual Macrophage Subset Suppresses Uterine Vascular Remodeling in Preeclampsia. Circulation. 2024 May 21;149(21):1670–88. doi:10.1161/CIRCULATIONAHA.123.064391 PubMed PMID: 38314577.

26. Sureshchandra S, Doratt BM, True H, Mendoza N, Rincon M, Marshall NE, et al. Multimodal profiling of term human decidua demonstrates immune adaptations with pregravid obesity. Cell Reports. 2023 Jul;42(7):112769. doi:10.1016/j.celrep.2023.112769

27. Dick SA, Wong A, Hamidzada H, Nejat S, Nechanitzky R, Vohra S, et al. Three tissue resident macrophage subsets coexist across organs with conserved origins and life cycles. Sci Immunol. 2022 Jan 7;7(67):eabf7777. doi:10.1126/sciimmunol.abf7777 PubMed PMID: 34995099.

28. Zhao J, Andreev I, Silva HM. Resident tissue macrophages: Key coordinators of tissue homeostasis beyond immunity. Sci Immunol. 2024 Apr 12;9(94):eadd1967. doi:10.1126/sciimmunol.add1967 PubMed PMID: 38608039.

29. Coleman M, Orvis A, Wu TY, Dacanay M, Merillat S, Ogle J, et al. A Broad Spectrum Chemokine Inhibitor Prevents Preterm Labor but Not Microbial Invasion of the Amniotic Cavity or Neonatal Morbidity in a Non-human Primate Model. Front Immunol. 2020;11:770. doi:10.3389/fimmu.2020.00770 PubMed PMID: 32425945; PubMed Central PMCID: PMC7203489.

30. Shynlova O, Boros-Rausch A, Farine T, Adams Waldorf KM, Dunk C, Lye SJ. Decidual Inflammation Drives Chemokine-Mediated Immune Infiltration Contributing to Term Labor. The Journal of Immunology. 2021 Oct 15;207(8):2015–26. doi:10.4049/jimmunol.2100493

31. Schultze JL. Reprogramming of macrophages--new opportunities for therapeutic targeting. Curr Opin Pharmacol. 2016 Feb;26:10–5. doi:10.1016/j.coph.2015.09.007 PubMed PMID: 26426255.

32. Samaniego R, Domínguez-Soto Á, Ratnam M, Matsuyama T, Sánchez-Mateos P, Corbí ÁL, et al. Folate Receptor β (FRβ) Expression in Tissue-Resident and Tumor-Associated Macrophages Associates with and Depends on the Expression of PU.1. Cells. 2020 Jun 10;9(6):1445. doi:10.3390/cells9061445 PubMed PMID: 32532019; PubMed Central PMCID: PMC7349916.

33. Cocker ATH, Whettlock EM, Browne B, Lai PF, Li JKH, Sivarajasingam SP, et al. Isolation of single cells from human uterus in the third trimester of pregnancy: myometrium, decidua, amnion and chorion. Oxford Open Immunology. 2022 Jun 11;3(1):iqac010. doi:10.1093/oxfimm/iqac010

34. Harshil Patel, Jonathan Manning, Phil Ewels, Maxime U Garcia, Alexander Peltzer, Rickard Hammarén, et al. nf-core/rnaseq: nf-core/rnaseq v3.27.0 - Vanadium Vulture [Internet]. Zenodo; 2026 [cited 2026 Oct 27]. Available from: https://zenodo.org/doi/10.5281/zenodo.1400710 doi:10.5281/ZENODO.1400710

35. Ge SX, Son EW, Yao R. iDEP: an integrated web application for differential expression and pathway analysis of RNA-Seq data. BMC Bioinformatics. 2018 Dec 19;19(1):534. doi:10.1186/s12859-018-2486-6 PubMed PMID: 30567491; PubMed Central PMCID: PMC6299935.

36. Kammerer J, Cirnu A, Williams T, Hasselmeier M, Nörpel M, Chen R, et al. Macro-based collagen quantification and segmentation in picrosirius red-stained heart sections using light microscopy. Biology Methods and Protocols. 2024 Jan 11;9(1):bpae027. doi:10.1093/biomethods/bpae027

37. Li M, Wu ZM, Yang H, Huang SJ. NFκB and JNK/MAPK activation mediates the production of major macrophage-or dendritic cell-recruiting chemokine in human first trimester decidual cells in response to proinflammatory stimuli. J Clin Endocrinol Metab. 2011 Aug;96(8):2502–11. doi:10.1210/jc.2011-0055 PubMed PMID: 21677045; PubMed Central PMCID: PMC3146787.

38. Huang SJ, Schatz F, Masch R, Rahman M, Buchwalder L, Niven-Fairchild T, et al. Regulation of chemokine production in response to pro-inflammatory cytokines in first trimester decidual cells. J Reprod Immunol. 2006 Dec;72(1–2):60–73. doi:10.1016/j.jri.2006.03.002 PubMed PMID: 16806486.

39. Hieshima K, Imai T, Baba M, Shoudai K, Ishizuka K, Nakagawa T, et al. A novel human CC chemokine PARC that is most homologous to macrophage-inflammatory protein-1 alpha/LD78 alpha and chemotactic for T lymphocytes, but not for monocytes. J Immunol. 1997 Aug 1;159(3):1140–9. PubMed PMID: 9233607.

40. Schraufstatter IU, Zhao M, Khaldoyanidi SK, Discipio RG. The chemokine CCL18 causes maturation of cultured monocytes to macrophages in the M2 spectrum. Immunology. 2012 Apr;135(4):287–98. doi:10.1111/j.1365-2567.2011.03541.x PubMed PMID: 22117697; PubMed Central PMCID: PMC3372745.

41. Gomez-Lopez N, StLouis D, Lehr MA, Sanchez-Rodriguez EN, Arenas-Hernandez M. Immune cells in term and preterm labor. Cell Mol Immunol. 2014 Nov;11(6):571–81. doi:10.1038/cmi.2014.46 PubMed PMID: 24954221; PubMed Central PMCID: PMC4220837.

42. Weiss A, Goldman S, Shalev E. The matrix metalloproteinases (MMPS) in the decidua and fetal membranes. Front Biosci. 2007 Jan 1;12:649–59. doi:10.2741/2089 PubMed PMID: 17127325.

43. Lash GE, Pitman H, Morgan HL, Innes BA, Agwu CN, Bulmer JN. Decidual macrophages: key regulators of vascular remodeling in human pregnancy. J Leukoc Biol. 2016 Aug;100(2):315–25. doi:10.1189/jlb.1A0815-351R PubMed PMID: 26819320.

44. Prins JR, Gomez-Lopez N, Robertson SA. Interleukin-6 in pregnancy and gestational disorders. J Reprod Immunol. 2012 Sep;95(1–2):1–14. doi:10.1016/j.jri.2012.05.004 PubMed PMID: 22819759.

45. Gschwandtner M, Derler R, Midwood KS. More Than Just Attractive: How CCL2 Influences Myeloid Cell Behavior Beyond Chemotaxis. Front Immunol. 2019;10:2759. doi:10.3389/fimmu.2019.02759 PubMed PMID: 31921102; PubMed Central PMCID: PMC6923224.

46. Rogers LM, Firestone K, Chinni R, Branco AC, Wrobleski K, Chou T, et al. Folate receptor beta drives NLRP3 inflammasome activation and pyroptosis in macrophages independent of folate binding. The Journal of Immunology. 2026 Apr 15;215(4):vkag051. doi:10.1093/jimmun/vkag051

47. Singh N, Herbert B, Sooranna G, Shah NM, Das A, Sooranna SR, et al. Is there an inflammatory stimulus to human term labour? PLoS One. 2021;16(8):e0256545. doi:10.1371/journal.pone.0256545 PubMed PMID: 34464407; PubMed Central PMCID: PMC8407546.

48. Menon R. Fetal inflammatory response at the fetomaternal interface: A requirement for labor at term and preterm. Immunol Rev. 2022 Jul;308(1):149–67. doi:10.1111/imr.13075 PubMed PMID: 35285967; PubMed Central PMCID: PMC9188997.

