## Supplementary information for "FOLR2 Identifies Functionally Distinct Decidual Macrophage Populations that Coordinate Immune and Tissue Remodelling During Human Labour"

**Supplementary Table 1**

| *Donor* | *Age* | *Parity* | *Gestation* | *BMI* | *Ethnicity* | *First language* | *Sex of baby* |
|---|---|---|---|---|---|---|---|
| *TNL_P1* | 39 | P1 | 39+0 | 21 | White, Any other white background | Unknown | Female |
| *TIL_P1* | 39 | P0 | 40+5 | 23.7 | White, Any other white background | Unknown | Female |
| *TIL_P2* | 31 | P0 | 41+6 | 18.4 | White, British | English | Female |
| *TNL_P3* | 32 | P1 | 39+1 | 22 | White, British | English | Female |
| *TIL_P3* | 30 | P0 | 40+5 | 20 | White, Any other white background | English | Male |
| *TIL_P4* | 35 | P0 | 40+2 | 18 | White, Any other white background | Lithuanian | Male |
| *TIL_P5* | 34 | P1 | 40+2 | 21 | White, Any other white background | English | Female |
| *TNL_P4* | 45 | P2 | 39+1 | 30 | Other, not stated | English | Female |
| *TNL_P5* | 43 | P1 | 39+1 | 20 | Mixed, Any other mixed background | Unknown | Male |
| *TNL_P6* | 28 | P1 | 39+0 | 18 | White, Any other white background | Unknown | Male |
| *TNL_P7* | 33 | P1 | 39+1 | 19 | White, Any other white background | Swedish | Male |
| *TNL_P8* | 34 | P0 | 39+3 | 25 | White, British | English | Female |
| *TNL_P9* | 40 | P5 | 37+5 | 20 | Other, any other ethnic group | English | Male |
| *TNL_P10* | 34 | P0 | 39+1 | 21 | White, British | English | Female |
| *TNL_P11* | 41 | P1 | 39+1 | 23 | White, Any other white background | Unknown | Female |
| *TIL_P6* | 32 | P1 | 40+4 | 23 | White, Any other white background | Moroccan | Female |

| | | | | | | | |
|---|---|---|---|---|---|---|---|
| *TIL_PTP1* | 29 | P0 | 33+0 | 22 | White, British | English | Female |
| *TNL_P12* | 42 | P1 | 39+3 | 23 | White, Any other white background | Romanian | Male |
| *TIL_P7* | 34 | P0 | 39+4 | 21 | White, Any other white background | English | Female |
| *T1_P1* | 36 | G1P0 | 7+ | 1st T | White, British | Unknown | N/A |
| *T1_P2* | 32 | G3P2 | 8+ | 1st T | Other, not stated | Unknown | N/A |
| *TIL_P8* | 40 | P2 | 40+2 | 21.7 | White, Irish | English | Male |
| *TIL_P9* | 38 | P3 | 41+0 | 22 | Black or Black British, African | Tigrinya | Female |
| *TIL_P10* | 26 | P1 | 39+6 | 18 | White, British | English | Male |
| *TNL_P13* | 37 | P1 | 39+0 | 30.8 | White, Any other white background | English | Male |
| *TNL_P14* | 35 | P2 | 38+4 | 22 | White, Any other white background | English | Female |
| *TNL_P15* | 33 | P0 | 39+2 | 26 | White, Any other white background | English | Female |
| *TNL_P16* | 44 | P0 | 39+0 | 24.9 | White, Any other white background | Portuguese | Male |
| *TNL_P17* | 37 | P0 | 39+5 | 19.3 | White, Any other white background | Portuguese | Female |
| *TNL_P18* | 39 | P1 | 38+5 | 25.8 | Other, any other ethnic group | Persian | Male |
| *TIL_P11* | 31 | P0 | 40+3 | 22 | White, Any other white background | Italian | Male |
| *TNL_P19* | 34 | P1 | 39+0 | 21.1 | White, British | English | Female |
| *TIL_P12* | 37 | P0 | 39+4 | 21.7 | Black or Black British, African | Zulu | Male |

| | | | | | | | |
|---|---|---|---|---|---|---|---|
| *TNL_P20* | 36 | P1 | 40+0 | 25.1 | White, British | English | Male |
| *TNL_P21* | 31 | P1 | 38+4 | 21.9 | Mixed - White and Black Caribbean | English | Male |
| *TIL_P13* | 35 | P0 | 40+2 | 18.5 | White, British | English | Male |
| *TNL_P22* | 36 | P1 | 39+5 | 21.2 | White, British | English | Female |
| *TNL_P23* | 30 | P0 | 39+0 | 29.3 | White, British | English | Male |
| *TNL_P24* | 32 | P0 | 40+0 | 24.1 | White, British | English | Male |
| *TNL_prac1* | 33 | P1 | 39+0 | 20.4 | Other, any other ethnic group | Portuguese | Female |
| *TNL_P25* | 36 | P2 | 38+6 | 22.3 | White, Any other white background | English | Female |
| *TNL_P26* | 37 | P1 | 39+5 | 20.7 | White, British | English | Female |
| *TeaL_P1* | 39 | P0 | 39+4 | 19.6 | Other, not stated | English | Female |
| *TNL_P27* | 35 | P0 | 39+2 | 26.4 | Black or Black British - African | English | Female |
| *TNL_P28* | 37 | P0 | 39+0 | 32.2 | White, Irish | English | Female |
| *TNL_P29* | 33 | P0 | 38+4 | 26 | White, British | English | Male |
| *TeaL_P2* | 39 | P0 | 38+4 | 23 | Other, not stated | English | Male |
| *TeaL_P3* | 28 | P1 | 38+6 | 19 | White, Any other White background | Italian | Female |
| *TIL_P14* | 31 | P0 | 40+1 | 21.8 | White, Any other White background | Spanish | Male |
| *TeaL_P4* | 40 | P2 | 38+3 | 25.6 | White, Any other White background | French | Female |
| *TIL_P15* | 34 | P0 | 40+4 | 21.5 | White, Any other White background | Italian | Female |

| | | | | | | | |
|---|---|---|---|---|---|---|---|
| *TIL_P16* | 38 | P1 | 41+2 | 25 | White, Any other White background | Spanish | Female |
| *TNL_P30* | 34 | P2 | 38+4 | 24.2 | White, Any other White background | English | Male |
| *TNL_P31* | 28 | P2 | 39+3 | 25.8 | Other, Any other ethnic group (middle eastern) | English | Female |
| *TeaL_P5* | 40 | P0 | 38+4 | 22.2 | Other, not stated | English | Male |
| *TNL_P32* | 37 | P1 | 38+0 | 25.2 | White, British | English | Female |
| *TNL_P33* | 42 | P1 | 39+1 | 29.3 | White, British | English | Male |
| *TNL_P34* | 32 | P0 | 39+2 | 23 | White, British | English | Female |
| *TNL_P35* | 32 | P1 | 38+2 | 21.4 | White, British | English | Female |
| *TNL_P36* | 30 | P2 | 37+4 | 29.3 | White British | English | Female |
| *TeaL_P6* | 37 | P1 | 39+1 | 20.7 | White, British | English | Male |
| *TeaL_P7* | 43 | P0 | 38+3 | 25.6 | White, any other white background | English | Male |
| *TNL_P37* | 32 | P0 | 40+2 | 24.2 | White, British | English | Female |
| *TNL_P38* | 34 | P1 | 39+0 | 22.2 | Other, any other ethnic group | Russian | Male |
| *TNL_P39* | 33 | P1 | 39+4 | 28.8 | White, British | English | Female |
| *TIL_P17* | 36 | P0+2 | 39+1 | 20.9 | White, British | English | Female |
| *TNL_P40* | 32 | G2 P0 | 39+5 | 22.5 | White, any other white background | English | Female |
| *TNL_P41* | 34 | P0 | 39+0 | 30.1 | Other, not stated | English | Male |
| *TNL_P42* | 35 | P2 | 38+3 | 36 | Black or Black British - African | Arabic | Female |

**Supplementary Figure 1**

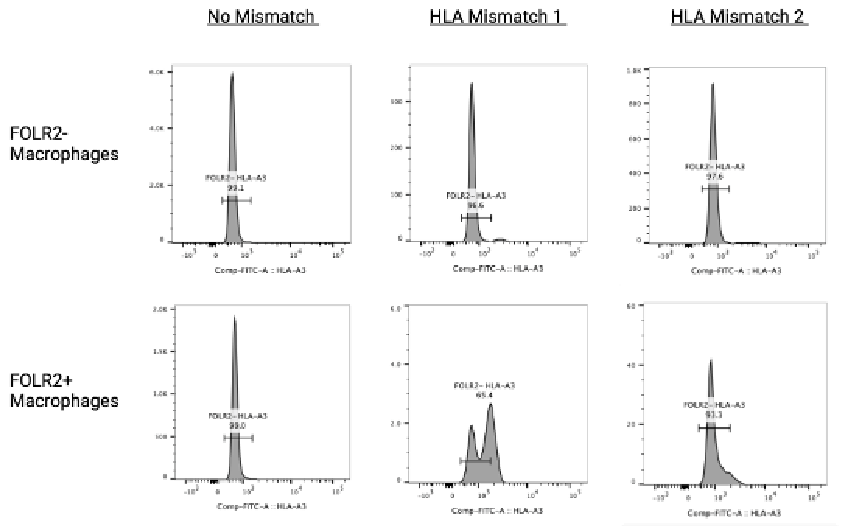


*Supplementary Figure 1. Identification of fetal-derived macrophages within the FOLR2+ compartment of the decidua basalis using HLA expression.*

Representative flow cytometry histograms showing HLA A3 expression within FOLR2- and FOLR2+ macrophage populations isolated from decidua basalis samples collected at term. Cells were gated on live, single, CD45+CD14+CD68+ macrophages prior to stratification by FOLR2 expression. FOLR2- macrophages were used to define the maternal HLA expression profile, while FOLR2+ macrophages were assessed for the presence of additional HLA A3 expression indicative of fetal origin. Examples include samples showing partial HLA mismatch within the FOLR2+ population, consistent with the presence of Hofbauer cells. Percentages indicate the proportion of cells within the indicated HLA-A3 maternal origin gate.

**Supplementary Figure 2**

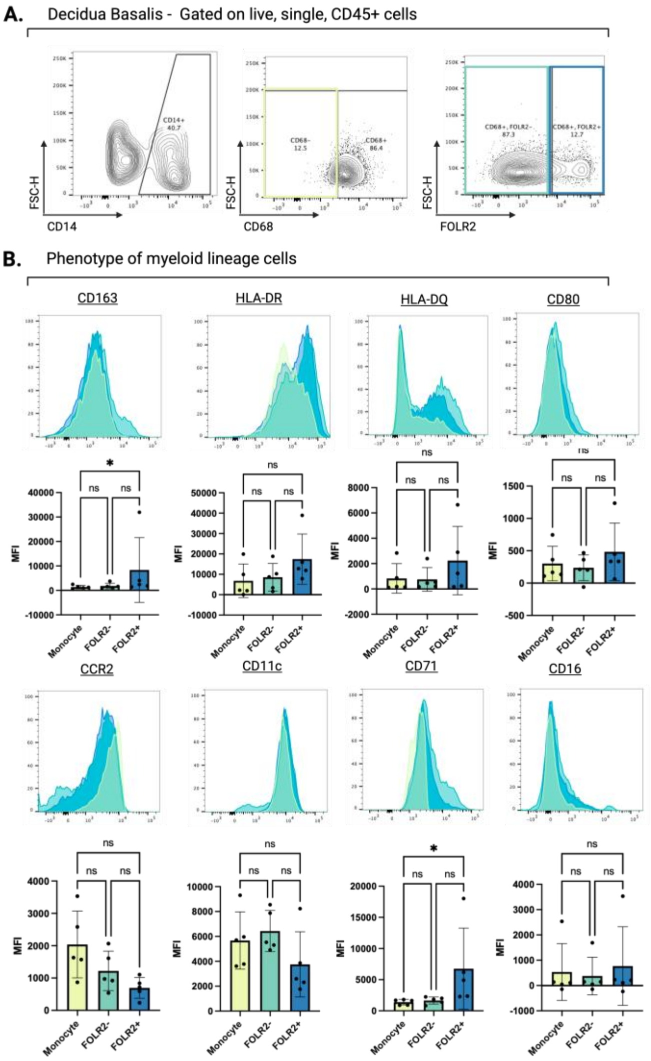


*Supplementary Figure 2. FOLR2 identifies two populations of macrophages cells in term decidua basalis*

Placentae with attached fetal membranes and decidua were collected from women delivering by elective caesarean section prior to the onset of labour at term. Immune cells were isolated from the decidua basalia and examined by flow cytometry. Immune cells were identified as live, single CD45+ cells. **A.** Myeloid lineage cells were identified as CD14+. Monocytes (yellow gate) were defined as CD68- and macrophages as CD68+. Within the macrophage population,

FOLR2- (green gate) and FOLR2+ (blue gate) subsets were defined. **B.** Expression of myeloid lineage markers by monocytes (yellow), FOLR2- (green) and FOLR2+ (blue) macrophages. Representative histograms and complete summary data for each marker examined is shown. Statistical comparisons were performed with Friedman's test and Dunn's test (non-normally distributed data: CD163, CD16, HLA-DR, CD71, and HLA-DQ) and RM one-way ANOVA with Tukey test (normally distributed data: CCR2, CD11c, and CD80) * $p < 0.05$; ** $p < 0.01$; *** $p < 0.001$; **** $p < 0.0001$.

**Commented [VM1]:** @Paterson, Emilie O Fill in the blanks and check I have the **** correct!

**Commented [PE2R1]:** used two diferent tests after normality testing, Friedmans or RM one-way. Feel free to change up the way i identified each one if you dont like it! Also - you are correct re. the *** !

**Supplementary Figure 3**

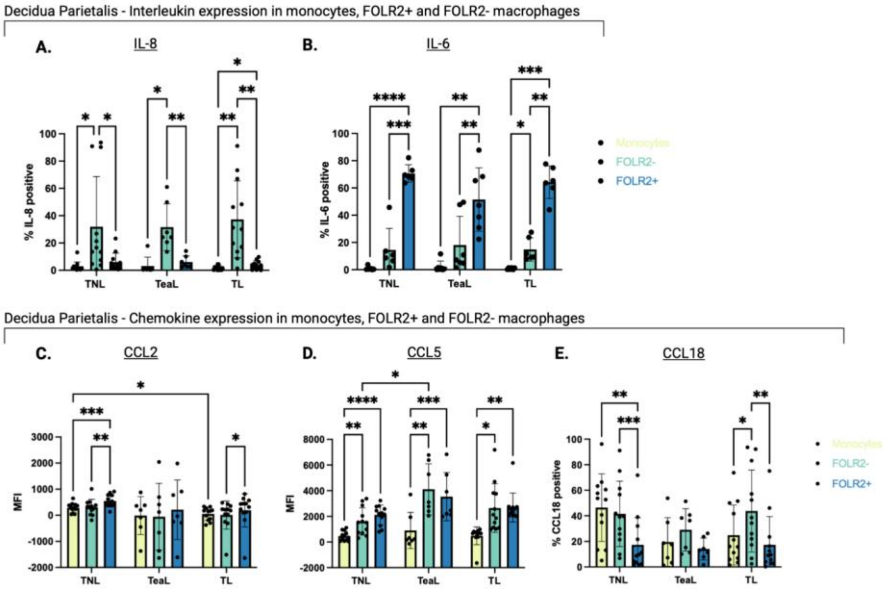


*Supplementary Figure 3. Intracellular flow cytometry results for monocytes, FOLR2- macrophages, and FOLR2+ macrophages in decidua parietalis across TNL, TeaL, and TL.* **A.** Interleukin -8 expression across TNL (n=12,), TeaL (n=7), and TL (n=12), respectively. **B.** Interleukin-6 across TNL (n=6), TeaL (n=7), and TL (n=6). **C.** CCL2 expression across TNL (n=12), TeaL (n=7), and TL (n=12). **D.** CCL5 expression across TNL (n=12), TeaL (n=7), and TL (n=12). **E.** CCL18 expression across TNL (n=12), TeaL (n=7), and TL (n=12). Two-way repeated-measures ANOVA with Tukey test. TNL, term not in labour; TEaL, term early labour; TL, term labour.

**Supplementary Figure 4**

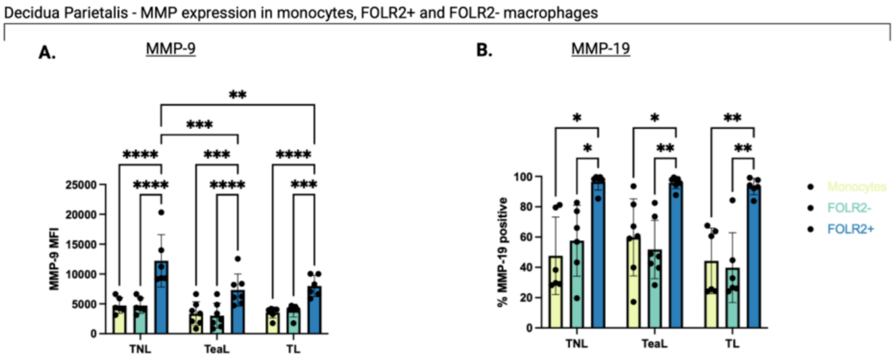


*Supplementary Figure 4. Expression of MMP-9 and MMP-19 in decidual monocytes, FOLR2- macrophages, and FOLR2+ macrophages across labour status.*

**A.** MFI of MMP-9 expression and **B.** percentage of MMP-19 positive cells across cell types in TNL, TeaL, and TL decidua parietalis. Each point represents an individual donor, and sample numbers for both panels were TNL n = 6, TeaL n = 7, and TL n = 6. Data were analysed using two-way repeated-measures ANOVA with Tukey's multiple comparisons test. TNL, term not in labour; TEaL, term early labour; TL, term labour.
